# Amino Acids Promote Mitochondrial-Derived Compartment Formation in Mammalian Cells

**DOI:** 10.1101/2020.12.23.424218

**Authors:** Max-Hinderk Schuler, Alyssa M. English, Leah VanderMeer, Janet M. Shaw, Adam L. Hughes

## Abstract

We recently identified a new cellular structure in yeast, called the Mitochondrial-Derived Compartment (MDC), that forms on mitochondria in response to amino acid excess. While emerging evidence supports an important function for MDCs in protecting cells from metabolic stress, whether this system exists beyond yeast remains unclear. Here, we show that MDCs are conserved in mammals, and like their yeast counterparts, are responsive to the intracellular amino acid content. Specifically, we find that inhibition of protein translation stimulates formation of dynamic, micron-sized compartments that associate with the mitochondrial network. These compartments are enriched for the carrier receptor Tomm70A and other select mitochondrial outer and inner membrane cargo, associate with the ER membrane, and require the conserved GTPase Miro1 for formation. Mammalian MDCs are responsive to changes in amino acid levels during translation inhibition, and are not activated by other common cellular stressors. Thus, MDCs represent an evolutionarily conserved nutrient-responsive mitochondrial remodeling system.

## INTRODUCTION

Mitochondria are double membrane-bound organelles that play essential roles in cellular energy production, amino acid homeostasis, and lipid metabolism, and their functional decline is associated with numerous age-related and metabolic disorders (Nunnari and Suomalainen, 2012). Cells utilize several systems to monitor, maintain and adjust mitochondrial function in response to various cellular stresses including protein aggregation, increased reactive oxygen species (ROS) formation, and loss of the mitochondrial membrane potential (ΔΨ) (Barbour and Turner, 2014). These quality control mechanisms include internal mitochondrial proteases that regulate turnover of proteins localized to the mitochondrial matrix, the inner membrane (IM), and the intermembrane space (IMS) (Quirós et al., 2015); the ubiquitin proteasome system, which controls clearance of proteins at the mitochondrial outer membrane (OM) (Karbowski and Youle, 2011); the mitochondrial unfolded protein response, which responds to protein aggregation stress within mitochondria (Shpilka and Haynes, 2018); mitochondrial fusion and fission, which integrate numerous signaling pathways to maintain balance of the mitochondrial network (Youle and van der Bliek, 2012); mitophagy, which promotes selective elimination of damaged mitochondria via Parkin-dependent and −independent autophagy (Pickles et al., 2018); and mitochondrial-derived vesicles (MDVs), which target damaged mitochondrial proteins from all sub-mitochondrial compartments for lysosomal degradation via small, sub-micron sized vesicles (70 – 150nm) in response to oxidative insults (Sugiura et al., 2014). Additionally, cells also modulate mitochondrial function, dynamics, and transport in response to numerous nutrient cues to maintain metabolic homeostasis during nutrient excess and caloric restriction (Liesa and Shirihai, 2013).

Among metabolic perturbations faced by cells, intracellular amino acid excess is a common defect in nutrient homeostasis that can alter mitochondrial function (Braun et al., 2015; Hughes et al., 2020; Li et al., 2017; Sun et al., 2016) and is associated with diabetes (Knebel et al., 2016; Newgard et al., 2009; Ruiz-Canela et al., 2018; Wang et al., 2011; Xu et al., 2013), cardio-vascular disease (Shah et al., 2010; Würtz et al., 2015) and a host of inborn errors of metabolism (Aliu et al., 2018). As with all nutrients, cells must tightly regulate amino acid levels by coordinating the rates of amino acid uptake, utilization and storage. To date, we understand a great deal about the signaling pathways and mechanisms that maintain cellular health during amino acid limitation (Efeyan et al., 2015; Rabinowitz and White, 2010). By contrast, how cells respond to toxic levels of amino acids that arise when systems that maintain amino acid homeostasis are disrupted is less well understood (Wellen and Thompson, 2010). We recently addressed this knowledge gap by identifying a new, nutrient-regulated cellular structure in yeast called the <u>M</u>itochondrial-<u>D</u>erived <u>C</u>ompartment (MDC) (Hughes et al., 2016; Schuler et al., 2020). MDCs are generated during cellular aging (Hughes et al., 2016) and in response to numerous insults that impair cellular amino acid homeostasis including defects in vacuolar amino acid compartmentation, inhibition of protein translation, and activation of amino acid acquisition programs in nutrient-replete conditions (Schuler et al., 2020). In response to amino acid elevation, MDCs form at contact sites between mitochondria and the ER in a manner dependent on the conserved mitochondrial outer membrane GTPase Gem1 (English et al., 2020). Upon formation, MDCs selectively sequester nutrient transporters of the SLC25A family and their associated import receptor Tom70 away from mitochondria, thereby providing cells with a mechanism to limit the abundance of nutrient transporters on mitochondria in response to amino acid excess (Schuler et al., 2020). Strikingly, MDC formation occurs simultaneously with removal of nutrient transporters from the plasma membrane via the multi-vesicular body (MVB) pathway and combined loss of MDC and MVB pathways renders cells sensitive to elevated amino acids. Thus, these pathways may operate as a coordinated network to protect cells from amino acid induced toxicity (Schuler et al., 2020). However, whether MDC formation is a conserved feature in higher eukaryotes remains unknown.

Here we set out to determine if MDCs form in mammalian cells and if core features of MDCs, i.e. substrate selectivity, association with the ER, dependence on the conserved GTPase Gem1 and regulation by amino acid abundance, are conserved. We show that inhibition of protein translation activates formation of micron-sized, Tomm70A-enriched, mitochondria-associated structures in cultured cells. Similar to yeast MDCs, these Tomm70A-enriched domains associate with the ER in mammalian cells and sequester select substrates from mitochondria, including a nutrient transporter of the SLC25A family. Formation of Tomm70A-enriched, MDC-like domains in mammalian cells requires the conserved GTPase Miro1, and is blocked in the absence of exogenous amino acids. Re-addition of single amino acids restores their formation in response to translation inhibition with glutamine being the most potent activator in cultured mammalian cells. By contrast, mitochondrial respiratory chain inhibition, increased ROS formation and protein aggregation stress do not activate the formation of MDC-like domains in mammals. Together, this work suggests that key features of MDC formation are conserved in mammalian cells and identifies a previously unrecognized, nutrient-regulated subcellular structure that forms on mammalian mitochondria during amino acid excess, a common perturbation in cellular nutrient homeostasis associated with numerous age-related and metabolic disorders.

## RESULTS

### Inhibition of protein translation activates formation of MDC-like domains in mammalian cells

We previously identified MDCs in the budding yeast *S. cerevisiae*, as large, Tom70-enriched, mitochondria-associated structures that exclude most other mitochondrial proteins, including the TIM23 translocase subunit Tim50, and form in response to numerous perturbations in amino acid homeostasis (Figure 1A). These perturbations included inhibition of vacuolar amino acid compartmentation via genetic or pharmacological impairment of the Vacuolar-H^+^-ATPase (V-ATPase), inhibition of protein biosynthesis, and activation of amino acid acquisition programs in nutrient-replete conditions by treating cells with the mechanistic target of Rapamycin (mTOR) inhibitor rapamycin (Schuler et al., 2020). To test if MDC formation is conserved in mammalian cells, we treated mouse embryonic fibroblasts (MEFs) stably expressing GFP-tagged Tomm70A and mRFP-tagged Timm50 with the V-ATPase inhibitor Concanamycin A (ConcA) (Dröse et al., 1993), Torin1, a specific inhibitor of mTOR signaling (Thoreen et al., 2009) or the protein translation inhibitor cycloheximide (CHX) (Schneider-Poetsch et al., 2010), and assayed for formation of Tomm70A-only containing structures that lack Timm50 by super-resolution live-cell imaging. As expected, Tomm70A and Timm50 co-localized throughout the tubular mitochondrial network in vehicle-treated cells (Figure 1B). Unlike yeast, treatment of MEFs with ConcA or Torin1 did not activate formation of any Tomm70A-only containing structures, despite efficiently blocking lysosomal acidification and mTOR signaling, as assayed via lysotracker staining and western blot analysis of phosphorylated mTOR substrates, respectively (Figure 1 Supplement 1A-H). This finding is consistent with recent reports showing that mammalian lysosomes do not store high amounts of amino acids like yeast vacuoles, and that mTOR inhibition blocks lysosomal amino acid egress in mammals without increasing whole-cell amino acid levels (Abu-Remaileh et al., 2017). By contrast, inhibition of protein translation with CHX induced formation of large, Tomm70A-enriched structures that lacked the IM protein Timm50 (Figure 1B). On average, these structures reached diameters of about one micron (900 nm ± 30 nm) (Figure 1B-C) which is similar to the size of late endosomes (Huotari and Helenius, 2011) and about ten times larger than the average reported size of mitochondrial-derived vesicles (Soubannier et al., 2012a). Time course analysis of the formation of these MDC-like domains revealed that they were visible as early as four hours after inhibition of protein translation with CHX (Figure 1D-E and Figure 1 Supplement 2). After eight hours, 86 ± 4% of all cells contained at least one MDC-like domain and most cells formed an average of 8.8 ± 0.5 per cell (Figure 1D-E). Interestingly, we observed at least one Tomm70A-enriched domain in about 10% of all untreated cells, indicating that these structures also form in the absence of CHX treatment, albeit at a low frequency (Figure 1D-E and Figure 1 Supplement 2).

**Figure 1.**
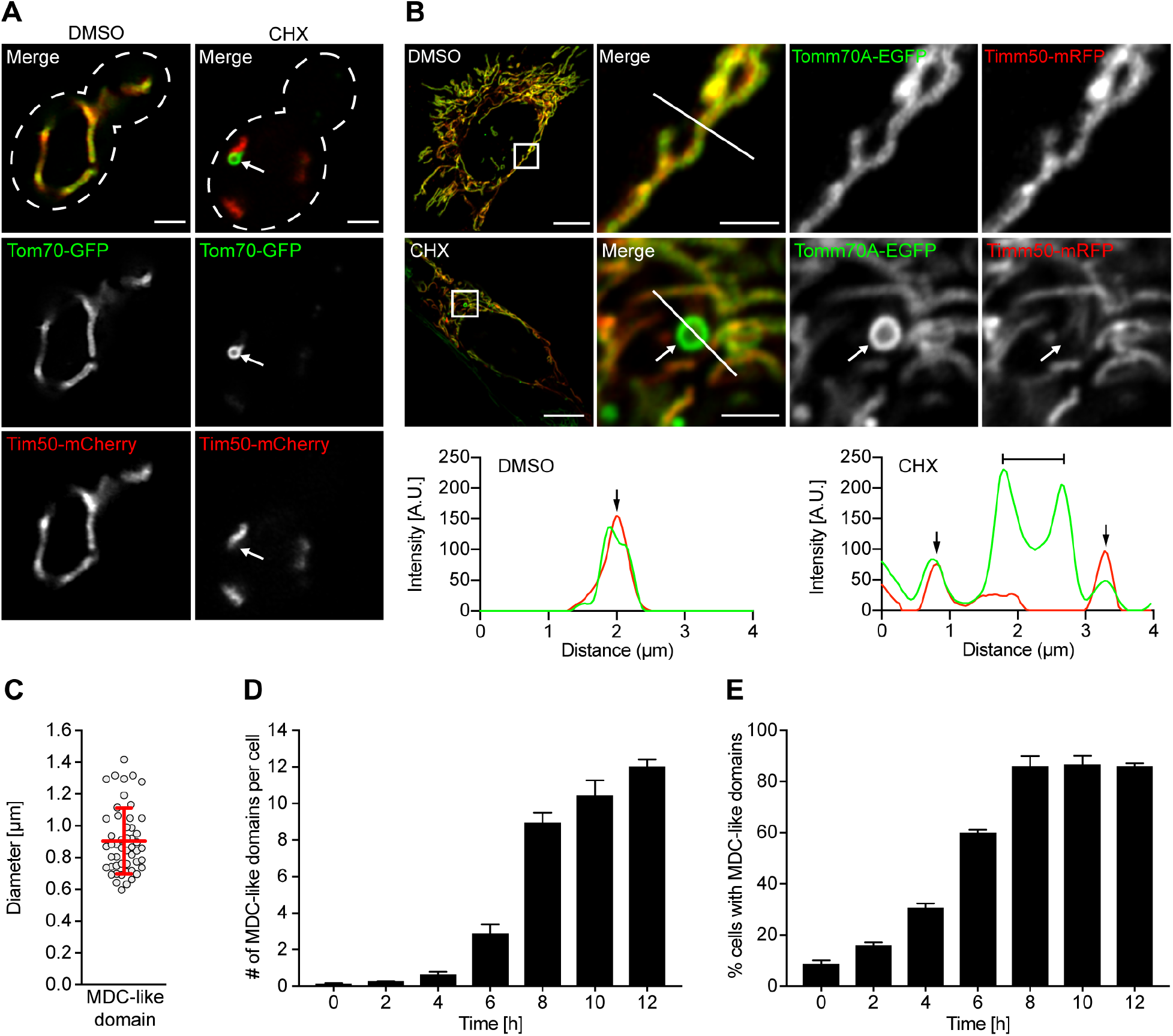
Translation inhibition activates formation of MDC-like domains in mammalian cells. (A) Images of cycloheximide (CHX)-induced MDCs in yeast expressing Tom70-GFP and Tim50-mCherry. White arrow marks MDC. Scale bar, 2 μm. (B) Images and line scan analysis of CHX-induced MDC-like domains in MEFs expressing Tomm70A-EGFP and Timm50-mRFP. White arrow marks MDC-like domain. White line marks fluorescence intensity profile position. Black arrow marks mitochondrial tubule and bracket marks MDC-like domain on line-scan analysis plot. Scale bars, 10 μm and 2 μm. (C) Scatter plot showing the diameter of CHX-induced MDC-like domains in MEFs. Error bars show mean ± SD of *n* = 50 MDCs from n = three experiments. (D, E) Quantification of CHX-induced MDC-like domain formation in MEFs over time. Error bars show mean ± SE from three replicates with *n* = 50 cells per replicate. (D) Number of MDC-like domains per cell. (E) Percentage of cells with at least one MDC-like domain.

### Mammalian MDC-like domains are cargo selective

We next sought to determine the substrate selectivity of this subcellular structure. In yeast, MDCs contain a distinct proteome, including members of the SLC25A family of IM nutrient transporters and their associated OM receptor Tom70. Other mitochondrial proteins localized to the matrix, the IM or the IMS are excluded from MDCs (Hughes et al., 2016). To test whether Tomm70A-enriched structures exhibit selectivity towards similar cargo in mammalian cells, we used indirect immunofluorescence (IIF) to localize various mitochondrial proteins upon translation inhibition. Similar to what occurs with yeast MDCs (Figure 2 Supplement 1A), we found that upon fixation and permeabilization, Tomm70A-enriched structures collapsed into puncta of reduced size with no visible lumen, suggesting that this cellular structure lacks factors that normally stabilize mitochondrial membranes (Figure 2 Supplement 1B). IIF revealed that Tomm70A-enriched domains excluded matrix proteins, including subunit beta of the mitochondrial ATP-Synthase (AtpB), the mitochondrial pyruvate dehydrogenase subunits E2/3 (PDH E2/3), and mitochondrial peroxiredoxin 3 (Prx3), though they were highly enriched for endogenous, untagged Tomm70A (Figure 2A and Figure 2 Supplement 1C). Similarly, Tomm70A-enriched domains also excluded IMS localized cytochrome c (Cyt C) (Figure 2B) and, like yeast MDCs, were not enriched for OM localized Tomm20 when compared to the mitochondrial tubule (Figure 2C). We also tested if MDC-like domains incorporate SLC25A nutrient transporters of the inner mitochondrial membrane, key constituents of yeast MDCs. Upon translation inhibition, Tomm70A-enriched, MDC-like domains selectively incorporated the GFP-tagged mitochondrial carrier protein SLC25A16 (Figure 2D), whereas another IM protein, Timm50, was excluded (Figure 1B and Figure 2 Supplement 1B). Additionally, MDC-like domains did not stain with the mitochondrial inner membrane potential-dependent dye tetramethylrhodamine methyl ester (TMRM) (Figure 2E), indicating that they lack a membrane potential, and are distinct structures compared to the mitochondrial tubules with which they associate. Finally, we found that CHX-induced formation and substrate selectivity of MDC-like domains were conserved across several commercially available cell lines including NIH 3T3 cells (mouse), Cos-7 cells (monkey) and 293T cells (human) (Figure 2 Supplement 2A-C). Together, these data indicate inhibition of protein translation activates formation of MDC-like structures in mammalian cells that have a cargo-selectivity comparable to that of yeast MDCs.

**Figure 2.**
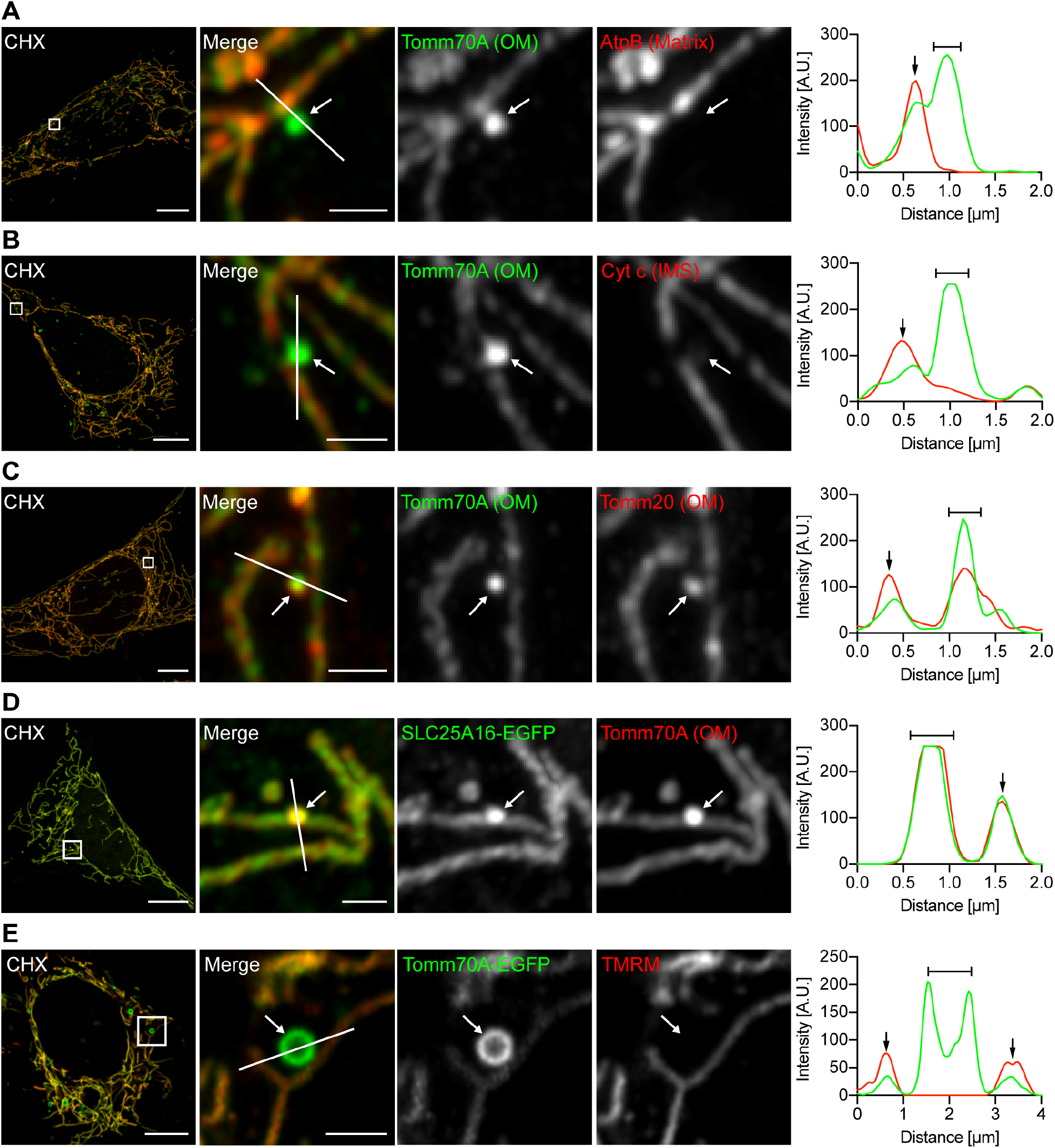
MDC-like domains incorporate select mitochondrial proteins. (A – D) Images and line scan analysis of cycloheximide (CHX) treated MEFs, fixed and stained for endogenous Tomm70A and the indicated mitochondrial protein. Outer membrane, OM; Intermembrane space, IMS; Inner membrane, IM. SLC25A16-EGFP was transiently expressed. (E) Images and line scan analysis of CHX treated MEFs expressing Tomm70A-EGFP and stained with the mitochondrial membrane potential-dependent dye TMRM. (A – E) White arrow marks MDC-like domain. White line marks fluorescence intensity profile position. Black arrow marks mitochondrial tubule and bracket marks MDC-like domain on line-scan analysis plot. Scale bars, 10 μm and 2 μm.

### MDC-like domains are dynamic and form from pre-existing mitochondria

We next sought to determine how MDC-like structures form in mammalian cells and gain a better understanding of their dynamics. To this end, we treated MEFs constitutively expressing Tomm70A-EGFP with CHX and analyzed formation of MDC-like domains by live-cell time-lapse imaging. In cultured mammalian cells, these compartments formed from mitochondria at distinct sites throughout the mitochondrial network (Figure 3A-B and Movies 1-2). Their formation was commonly preceded by a local expansion of the mitochondrial diameter followed by a fast collapse and accumulation of Tomm70A in the MDC-like domain (Figure 3A-B). Upon formation, most Tomm70A-enriched structures remained associated with the mitochondrial network for extended periods of time (Movies 1-2). We also occasionally observed release of MDC-like domains from mitochondria and subsequent transient interactions with the network, including docking of the MDC-like structure to mitochondria and movement along mitochondrial tubules (Figure 3C and Movie 3). In summary, MDC-like domains originate from pre-existing mitochondria and display dynamics comparable to other cellular organelles.

**Figure 3.**
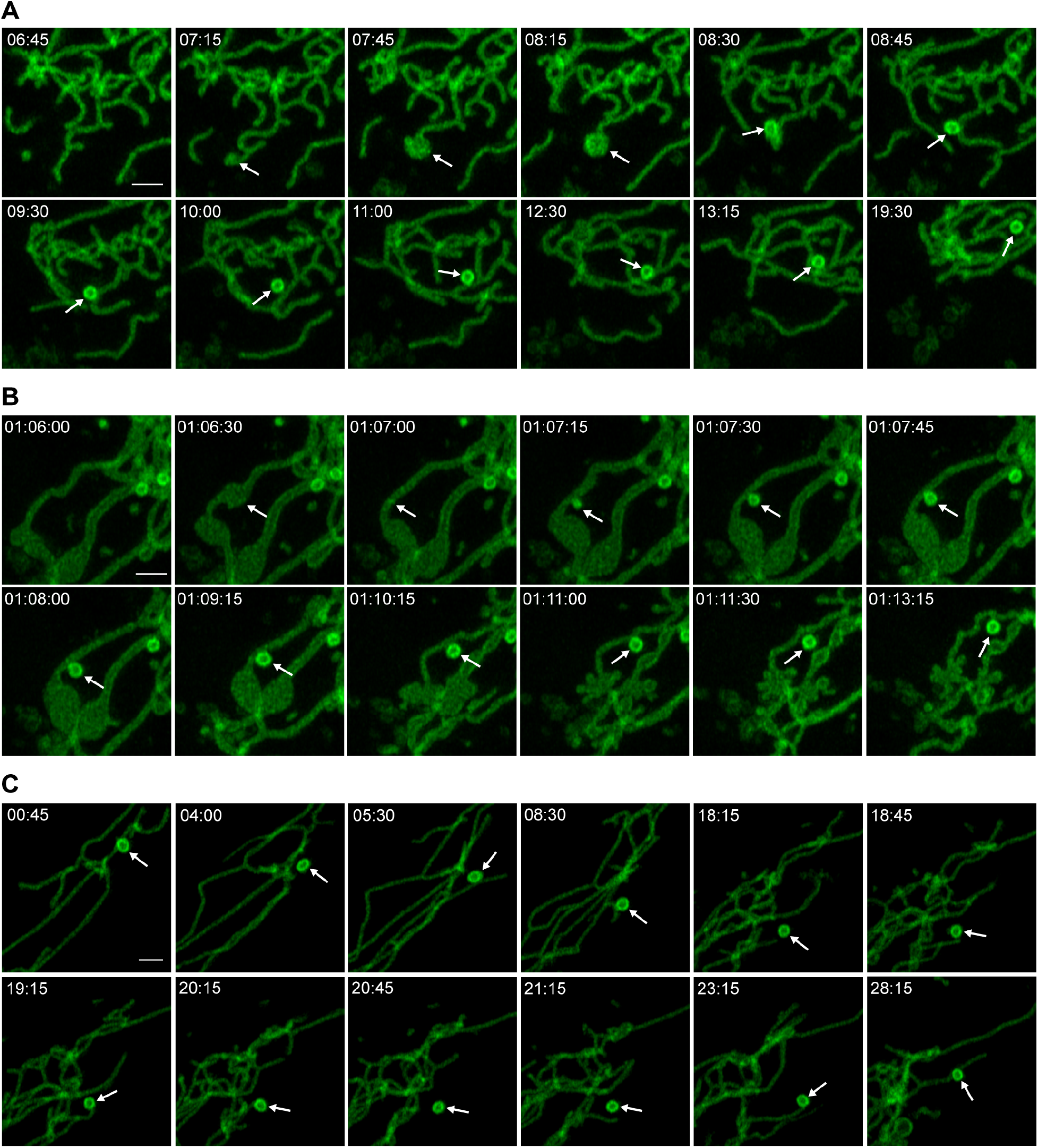
MDC-like domains form from mitochondria and exhibit dynamic behaviors. (A, B) Time-lapse images of cycloheximide (CHX)-induced MDC-like domain formation in MEFs expressing Tomm70A-EGFP. MDC-like domain formation is commonly preceded by a local expansion of the mitochondrial tubule diameter. (C) Time-lapse images of MDC-like domain dynamics in MEFs expressing Tomm70A-EGFP. MDC-like domains are released from mitochondria and can subsequently re-associate with the local network. (A – C) Arrowhead marks MDC-like domain. Scale bar, 2 μm.

### MDC-like domains associate with the ER in mammalian cells

Because of their dynamic, organelle-like behavior, we next tested if MDC-like domains co-localize with other cellular organelles including the ER, lysosomes, endosomes and peroxisomes. We first treated MEFs expressing the ER marker BFP-KDEL and Tomm70A-EGFP with CHX and found that 99% of all MDC-like structures were associated with ER tubules (Figure 4A, C and Figure 4 Supplement 1A). Interestingly, we observed that MDC-like structures that had been released from the mitochondrial network remained associated with the ER, suggesting a stable interaction between these two cellular compartments (Figure 4A and Figure 4 Supplement 1A). In line with this observation, we recently found that MDCs form at mitochondria-ER contact sites in yeast and remain associated with the ER over extended periods (English et al., 2020). By contrast, only about 20% of all Tomm70A-enriched domains associated with lysosomes (Figure 4B-C and Figure 4 Supplement 1B) and a very small fraction associated with EEA1-TagRFP labeled endosomes or Pex14 labeled peroxisomes (Figure 4B-C and Figure 4 Supplement 1C-D). Together these data suggest that similar to their yeast counterparts, MDC-like domains stably associate with the ER in mammals whereas interactions with other organelles are likely more transient.

**Figure 4.**
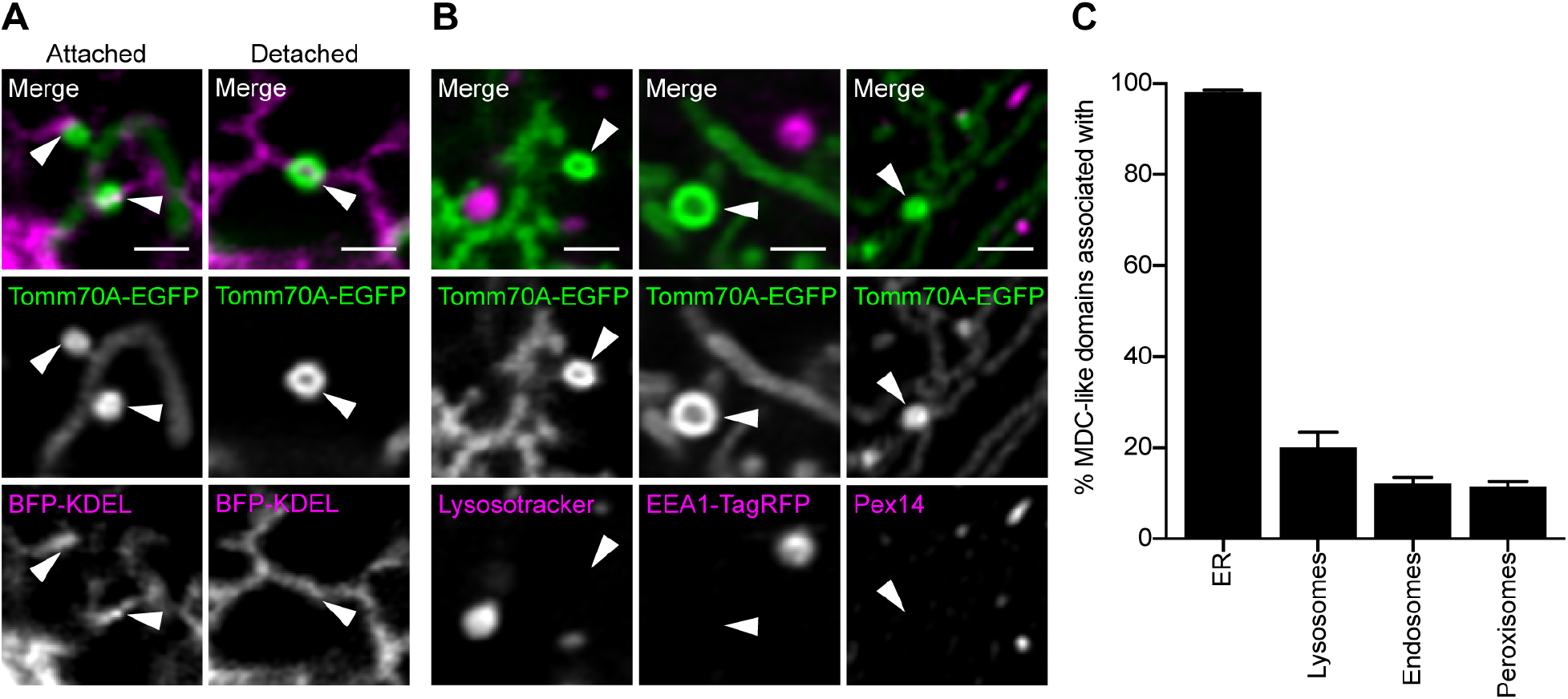
MDC-like domains associate with the ER in mammalian cells. (A) Images of cycloheximide (CHX)-induced MDC-like domains in MEFs expressing Tomm70-A-EGFP and the ER marker BFP-KDEL. MDC-like domains associate with the ER, both when attached to or detached from the mitochondrial network. Arrowhead marks MDC-like domain. Scale bar, 2 μm. (B) Images of CHX-induced MDC-like domains in MEFs expressing Tomm70-A-EGFP. Lysosomes were stained with the pH-sensitive dye Lysotracker Red. Endosomes were labeled by expression of EEA1-TagRFP. Peroxisomes were visualized upon fixation, permeabilization and staining for the peroxisomal protein Pex14. Arrowhead marks MDC-like domain. Scale bar, 2 μm. (C) Quantification of MDC-like domain association with the ER, lysosomes, endosomes or peroxisomes. Error bars show mean ± SE from three replicates with *n* = 50 MDC-like domains per replicate.

### The conserved GTPase Miro1 is required for formation of MDC-like domains in mammalian cells

In yeast, mitochondria are linked to the ER via the ER-mitochondria encounter structure (ERMES) (Kornmann et al., 2011, 2009) and we recently showed that ERMES and its associated GTPase Gem1 are required for MDC formation (English et al., 2020). We thus tested if the Gem1 orthologs Miro1 and Miro2 (Fransson et al., 2003) are required for formation of the Tom70A-enriched structures we identified in mammalian cells. To this end, we quantified CHX-induced formation of MDC-like domains in MEFs isolated from *Miro1^-/-^* (Nguyen et al., 2014) and *Miro2^-/-^* embryos (López-Doménech et al., 2016) and their corresponding littermate controls. In *Miro1^+/+^* MEFs, inhibition of protein translation induced an average of 8.9 ± 1.3 MDC-like domains per cell in 88.7 ± 1.8 % of all cells, whereas formation of these Tomm70A-enriched structures was largely blocked upon deletion of *MIRO1* (0.3 ± 0.1 MDC-like domains per cell in 11.3 ± 1.3 % of all cells) (Figure 5A-B). By contrast, deletion of *MIRO2* only mildly affected CHX induced formation of MDC-like structures in MEFs (Figure 5C-D). Thus, at least in MEFs, Miro1 plays a critical role for formation of Tomm70A-enriched, MDC-like domains in response to translation inhibition, while Miro2 seems mostly dispensable. These data indicate that the conserved GTPase Miro1, a key factor required for MDC formation in yeast also promotes formation of MDC-like domains in mammalian cells.

**Figure 5.**
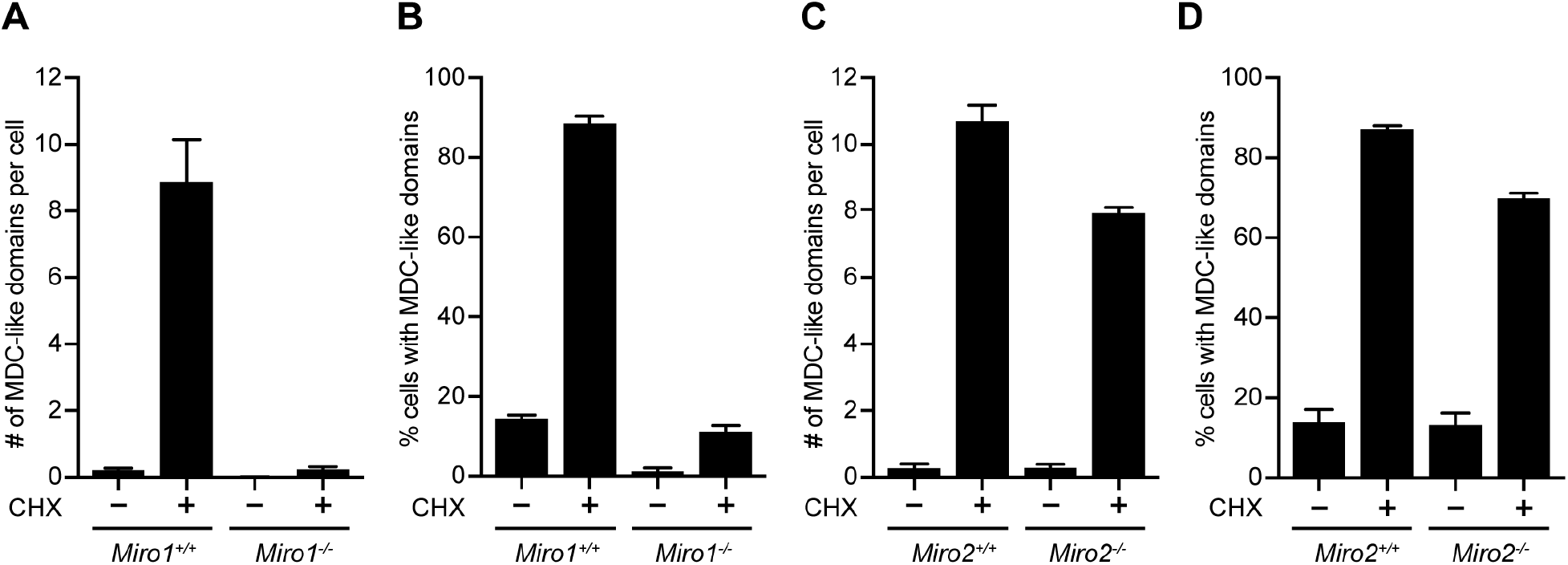
Miro1 is required for formation of MDC-like domains in mammalian cells. (A, B) Quantification of cycloheximide (CHX)-induced MDC-like domain formation in *Miro1*^+/+^ and *Miro1*^-/-^ MEFs. (C, D) Quantification of CHX-induced MDC-like domain formation in *Miro2*^+/+^ and *Miro2*^-/-^ MEFs. (A, C) Number of MDC-like domains per cell. (B, D) Percentage of cells with at least one MDC-like domain. (A – D) Error bars show mean ± SE from three replicates with *n* = 50 cells per replicate.

### High levels of amino acids are required for formation of MDC-like domains in mammals

Lastly, we sought to determine whether amino acid levels play a role in activating MDC-like domain formation in response to translation inhibition. In a recent study, we found that yeast cells form MDCs in response to an overabundance of intracellular amino acids and showed that impairment of protein translation activated MDC formation by increasing intracellular amino acid levels (Schuler et al., 2020). We thus tested if formation of mammalian MDC-like structures in response to CHX is blocked in the absence of exogenous amino acids. To this end, we cultured cells in media lacking all amino acids and quantified formation of Tomm70A-enriched, MDC-like structures in response to translation inhibition by CHX. In normal culture media, inhibition of protein biosynthesis increased intracellular amino acid levels as previously reported (Figure 6 Supplement 1A), and robustly activated formation of Tomm70A-enriched domains in multiple cell lines including MEFs (Figure 6A-B and Figure 6 Supplement 1B-C), Cos-7 (Figure 6 Supplement 1D-E) and 293T cells (Figure 6 Supplement 1F-G). By contrast, formation of MDC-like structures was blocked in all tested cell lines in the absence of exogenous amino acids, indicating that high amino acids are required for their formation in response to translation inhibition (Figure 6A-B and Figure 6 Supplemental 1B-G).

**Figure 6.**
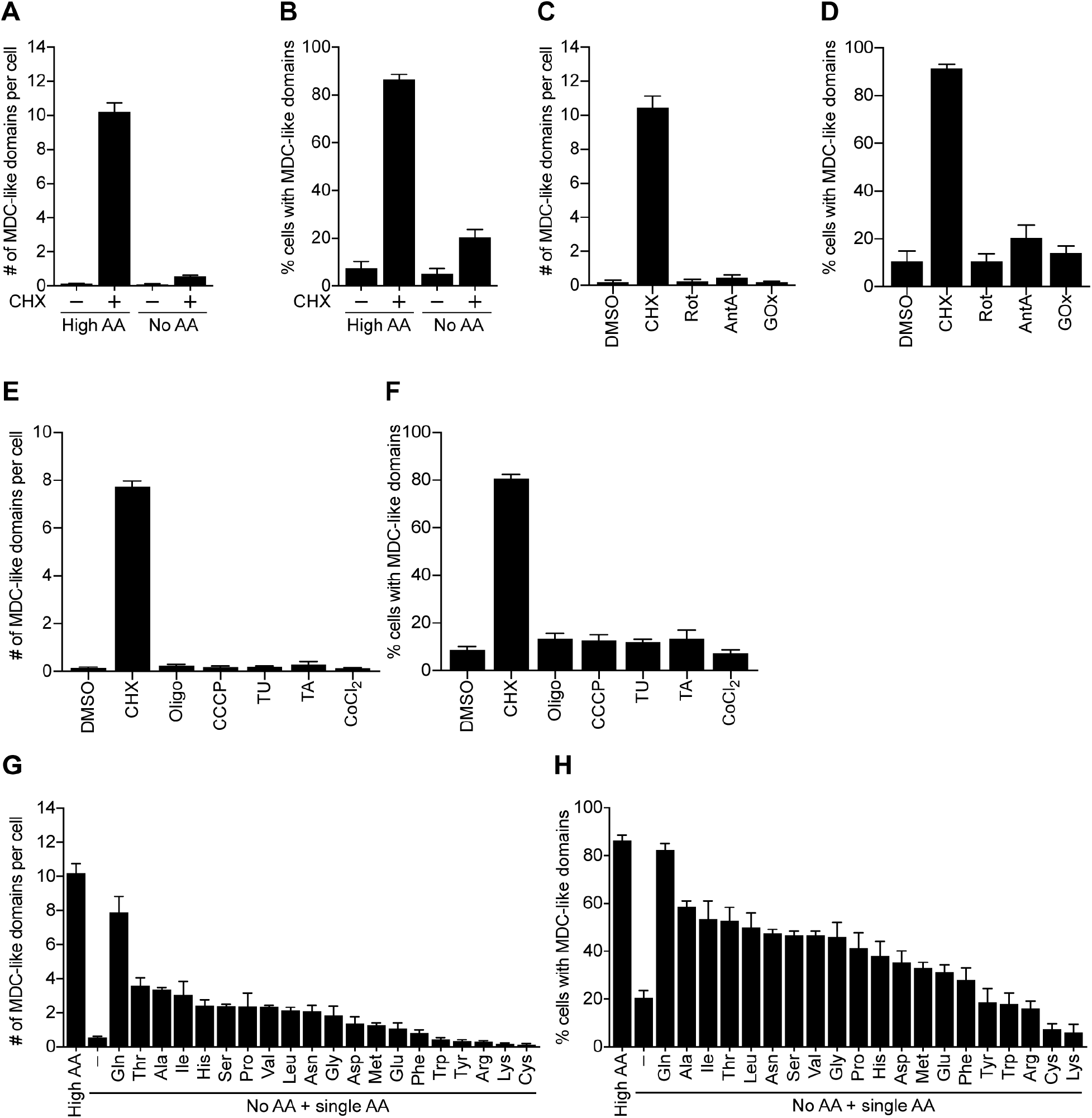
Amino acid excess stimulates formation of MDC-like domains in mammalian cells. (A, B) Quantification of cycloheximide (CHX)-induced MDC-like domain formation in media containing high levels of amino acids or amino acid free media. (C, D) Quantification of MDC-like domain formation in response to rotenone (Rot), antimycin A (AntA) and glucose oxidase (GOx). (E, F) Quantification of MDC-like domain formation in response to inhibition of ATP production with oligomycin (Oligo), depolarization of mitochondria with CCCP, or induction of ER-stress with tunicamycin (TU), thapsigargin (TA) or CoCl2. (G, H) Quantification of CHX-induced MDC-like domain formation upon re-addition of single amino acids to amino acid free media. (A, C, E, G) Number of MDC-like domains per cell. (B, D, F, H) Percentage of cells with at least one MDC-like domain. (A – H) Error bars show mean ± SE from three replicates with *n* = 50 cells per replicate.

We recently showed that increased cellular amino acids, specifically cysteine, impair mitochondrial respiratory chain function by limiting cellular iron bioavailability through an ROS based mechanism (Hughes et al., 2020). Notably, numerous mitochondrial quality control pathways, including mitophagy (Schofield and Schafer, 2020) and mitochondrial-derived vesicles (Soubannier et al., 2012a, 2012b), respond to elevated cellular ROS. We thus considered that ROS may activate MDC-like structures downstream of amino acid elevation in mammalian cells. To test this possibility, we quantified formation of MDC-like domains in response to various ROS-inducing agents including the mitochondrial respiratory chain complex I inhibitor rotenone (Rot), the complex III inhibitor antimycin A (AntA), and Glucose Oxidase (GOx), a source of cytosolic H2O2. Consistent with previous reports, AntA and Rot increased mitochondrial peroxiredoxin 3 (Prx3) dimerization, indicating an increase in mitochondrial ROS production, whereas GOx induced hyper oxidation of cytosolic Prx2 (Cox et al., 2009), indicating increased ROS production in the cytoplasm (Figure 6 Supplement 2A). However, neither of these treatments activated formation of Tomm70A-enriched structures (Figure 6C-D) indicating that, unlike other mitochondrial quality control mechanisms, MDC-like domains are not activated in response to increased ROS. In support of this idea, inhibition of protein translation did not affect the redox balance of cytosolic Prx2 or mitochondrial Prx3 (Figure 6 Supplement 2A). Moreover, culturing cells in the presence of galactose, which promotes mitochondrial ROS formation through increased mitochondrial respiration (Benard et al., 2007), or supplementing media with the antioxidant n-acetylcysteine (Sun, 2010), did not activate or block formation of MDC-like structures, respectively (Figure 6 Supplement 2B-C). Lastly, inhibition of mitochondrial ATP production with oligomycin, uncoupling of the mitochondrial membrane potential with CCCP or activation of the ER-stress response with tunicamycin (TU), thapsigargin (TA) or CoCl2, all failed to stimulate formation of MDC-like domains (Figure 6E-F).

### Single amino acids promote MDC-like domain formation during translation inhibition in mammalian cells

Our data indicate that MDC-like domains are formed in response to intracellular amino acid elevation during translation inhibition, and are unresponsive to other cellular stresses. We thus determined which amino acids are required to activate MDC-like structures in MEFs. Re-addition of single amino acids to media lacking all other amino acids robustly activated CHX-induced formation of Tom70A-enriched structures, indicating that their formation was directly responsive to the supply of exogenous amino acids (Figure 6G-H). Strikingly, glutamine, the most abundant amino acid in sera and tissue culture media, almost completely restored formation of Tomm70A-enriched structures in response to CHX, followed by several other amino acids that drive MDC formation in yeast, including alanine, threonine, histidine and several branched-chain amino acids (BCAAs) (Figure 6G-H). By contrast, re-addition of cysteine, which impairs cellular iron homeostasis through an ROS-based mechanism in yeast (Hughes et al., 2020), did not promote formation of MDC-like domains (Figure 6G-H). Together, these data suggest that MDC-like domains are formed in response to intracellular amino acid overabundance during translation inhibition.

## DISCUSSION

We previously showed that intracellular amino acid overabundance activates formation of a newly described, mitochondria-associated sub-compartment in yeast, the MDC (Hughes et al., 2016; Schuler et al., 2020). Here we present evidence that key features of this previously unrecognized cellular structure are conserved in mammalian cells (Figure 7). First, we show that mammalian MDC-like domains are large and dynamic structures that form from mitochondria and are in contact with the ER. Second, we demonstrate that mammalian MDC-like structures are selectively enriched for Tomm70A and that these domains can incorporate a nutrient carrier of the SLC25A family. By contrast, other IM proteins as well as proteins localized to the mitochondrial IMS and the mitochondrial matrix are excluded from these Tomm70A-enriched compartments. Third, we show that formation of MDC-like structures in mammalian cells requires the conserved GTPase Miro1. Lastly, our data indicate that MDC-like domains form in response to amino acid overabundance but are not responsive to other cellular stresses such as mitochondrial dysfunction, increased ROS, or ER-stress. Based on this collection of observations, we propose that these mammalian structures are indeed the counterpart of yeast MDCs, suggesting these unique mitochondrial-associated domains are evolutionarily conserved in mammals.

**Figure 7.**
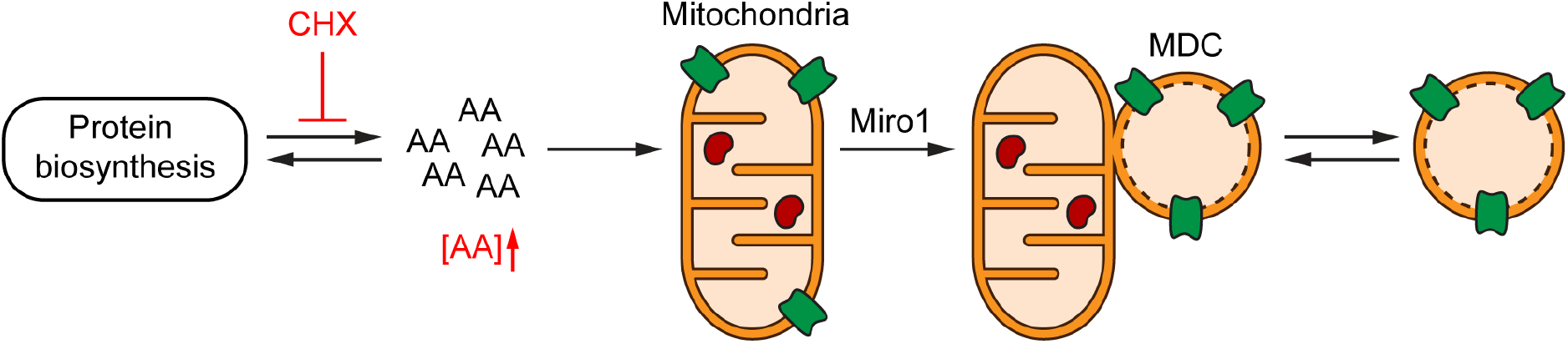
Translation inhibition activates amino acid- and Miro1-dependent remodeling of mitochondria in mammals. Model of MDC formation in mammalian cells. In response to amino acid elevation, MDCs sequester select proteins (green) from mammalian mitochondria, while excluding most other mitochondrial proteins (red).

With the discovery of MDCs in mammals, a key question that arises is whether or not these structures are related to the recently characterized mitochondrial-derived vesicles. While we cannot exclude the possibility that these two pathways may ultimately be related, at this point, data from studies of both systems suggest they are likely distinct. MDVs are vesicular structures, often ranging in size from 70 – 150nm, that move cargo from all mitochondrial sub-compartments to other organelles, including lysosomes and peroxisomes (Neuspiel et al., 2008; Soubannier et al., 2012a). MDV formation is commonly activated by oxidative stress, and their biogenesis is not known to occur at ER-mitochondrial contact sites. MDCs on the other hand are larger cellular structures that reach sizes approaching 1 μm. While they are cargo selective, they do not appear to traffic from the mitochondria to other organelles. Instead, they form at and dynamically associate with mitochondria-ER contact sites, where they selectively sequester the carrier import receptor Tomm70A and its clients in both yeast and mammals. Finally, MDC formation is unresponsive to ROS stress and requires Gem1 in yeast and Miro1 in mammalian cells, and MDC release from mitochondria relies on the mitochondrial fission GTPase Dnm1/Drp1 in yeast, which does not promote MDV fission in mammals. Thus, MDCs appear to differ from MDVs with regard to size, substrate selectivity, regulation, and formation mechanism. Moving forward, it will be interesting to investigate the interplay between these systems and their role in regulating mitochondrial and cellular physiology.

The establishment of this new conserved structure also raises important questions for future studies. Chief amongst them is whether MDC formation provides cells with a mechanism to fine-tune mitochondrial metabolism in response to amino acid excess, or if the selective enrichment of Tom70 and its clients in MDCs serves a different cellular purpose. Moreover, how amino acid excess is sensed and relayed to MDC formation machinery is unknown. In yeast, branched-chain amino acids (BCAAs) and their catabolites activate MDC formation (Schuler et al., 2020), whereas glutamine most potently activates MDC formation in cultured mammalian cells. As both BCAA transamination and glutamine catabolism are linked to the tricarboxylic acid (TCA) cycle through glutamate, α-ketoglutarate and acetyl-CoA (Cluntun et al., 2017; Neinast et al., 2019), it is possible that a common downstream metabolite serves as a signaling intermediate to activate MDC formation. Alternatively, amino acid signal may also be directly relayed to MDC formation machinery through specialized amino acid sensing systems. In either case, we anticipate that clues regarding the sensing mechanism may come from studies of other nutrient responsive systems including the ESCRT/MVB system (Risinger et al., 2006; Rubio-Texeira and Kaiser, 2006), mTOR signaling (González and Hall, 2017; Wolfson and Sabatini, 2017), and a multitude of conserved cellular energy and metabolite sensors (Chantranupong et al., 2015; Ljungdahl and Daignan-Fornier, 2012).

We showed here that the conserved GTPase Miro1 is required for MDC formation in cultured cells. In yeast, the Miro1 ortholog Gem1 is a regulatory subunit of the ERMES complex (Kornmann et al., 2011), which facilitates lipid exchange between mitochondria and the ER (Kawano et al., 2018; Kojima et al., 2016). However, ERMES is not completely conserved in mammalian cells (Herrera-Cruz and Simmen, 2017), and lipid transport between mitochondria and the ER seems to be dispensable for MDC formation in yeast (English et al., 2020), raising the possibility that Miro1 and Gem1 may play a more direct role in MDC formation. One possible scenario is that Gem1 in yeast and Miro1 in mammals promote MDC formation by connecting mitochondria to the cytoskeleton. In support of this idea, Miro1 links mitochondria to microtubules and actin filaments in mammalian cells (Fransson et al., 2006; Guo et al., 2005; López-Doménech et al., 2018; Nguyen et al., 2014) and cytoskeletal forces play important roles in numerous other cellular processes that require cargo sorting and membrane re-organization (Granger et al., 2014; Grant and Donaldson, 2009; Gurel et al., 2014). Notably, while deletion of *MIRO1* confines the mitochondrial network to the perinuclear area (Nguyen et al., 2014) and blocks MDC formation in MEFs, deletion of *MIRO2* does not impair the intracellular distribution of mitochondria in this cell type (López-Doménech et al., 2016) and has little to no effect on MDC formation. Alternatively, Miro proteins have recently been shown to regulate mitochondria-ER contacts in mammalian cells (Modi et al., 2019) and contacts between the two organelles may provide unknown factors required for MDC formation. Moving forward, it will be important to determine the mechanism by which Gem1 and Miro1 promote MDC formation in yeast and mammals, how high levels of amino acids are sensed to activate MDC formation, and under what conditions and in which tissues MDCs form *in vivo*. Answers to these key questions will provide insights into how formation of this previously unrecognized, conserved cellular structure contributes to cellular homeostasis during amino acid excess.

## Acknowledgements

We thank members of the A.L.H. and J.M.S. laboratories for discussion and manuscript comments. Metabolomics analysis was performed at the University of Utah Metabolomics Core directed by J. Cox and supported by National Institutes of Health (NIH) grants 1S10OD016232-01, 1S10OD021505-01 and 1U54DK110858-01. Research was supported by NIH grants GM119694 and AG061376 (A.L.H.), NIH T32GM007464 (A.M.E.), NIH GM53466 and GM84970 (J.M.S.), AHA 18PRE33960427 (M.H.S.) and the Howard Hughes Medical Institute (J.M.S.). A.L.H. was further supported by a Searle Scholars Award and Glenn Foundation for Medical Research Award.

## Author Contributions

All authors conceived aspects of the project, designed experiments, and discussed and analyzed results. M.H.S, A.M.L. and L.V. conducted experiments. M.H.S. and A.L.H wrote and edited the manuscript.

## Materials and Methods

### Yeast Strains

*Wild-type* yeast strain AHY4706 has been previously described (Schuler et al., 2020) and is a derivative of *Saccharomyces cerevisiae* S288c (BY) (Brachmann et al., 1998) expressing fluorescently tagged *TOM70* and *TIM50* from their native loci. AHY4706, was rendered prototrophic with pHLUM, a yeast plasmid expressing multiple auxotrophic marker genes from their endogenous promoters (Mülleder et al., 2012), to prevent complications caused by amino acid auxotrophies in the BY strain background.

### Plasmids

To generate pLenti-mTomm70A-EGFP the open reading frame of mouse Tomm70A (GenScript OMu13526) was amplified via PCR and inserted into pcDNA-EGFP introducing a ten amino acid long linker between mTomm70A and EGFP (GGSGDPPVAT). mTomm70A-EGFP was then transferred into pQCXIN (Clontech #631514) to generate pLenti-mTomm70A-EGFP. To generate pLenti-mTimm50-mRFP the open reading frame of mouse Timm50 (GenScript OMu13400) was amplified via PCR and inserted into pcDNA-mRFP introducing a ten amino acid long linker between mTimm50 and mRFP (GGSGDPPVAT). The internal EcoRI site was disrupted by introducing at silent mutation at nt1283 (G to A) using Q5 site directed mutagenesis (NEB) and mTimm50-mRFP was cloned in to pQCXIN (Clontech #631514) to generate pLenti-mTimm50-mRFP. pcDNA6.2-hSLC25A16-EmGFP was generated by cloning human SLC25A16 cDNA, obtained from the human ORF collection (Rual et al., 2005), into pcDNA6.2/C-EmGFP-DEST using Gateway LR Clonase II (ThermoFisher). TagBFP-C-KDEL (# 49150), TagRFP-T-EEA1 (# 42635) and pHLUM (# 40276) were obtained from Addgene (# 49150 and # 42635) and have been described previously (Friedman et al., 2011; Mülleder et al., 2012; Navaroli et al., 2012).

### Yeast Cell Culture and MDC Assays

For yeast MDC assays, cells were grown exponentially for 15 hours at 30°C to a maximum density of 6×10^6^ cells/ml in media containing high amino acids (1% yeast extract, 2% peptone, 0.005% adenine, 2% glucose). This period of overnight log-phase growth was carried out to ensure vacuolar and mitochondrial uniformity across the cell population and is essential for consistent MDC activation. To induce MDC formation, cycloheximide (CHX) (Sigma-Aldrich; C1988) was added to overnight-log phase cultures at a final concentration of 10 μg/ml for two hours. After incubation, cells were harvested by centrifugation and either resuspended in imaging buffer (5% w/v glucose, 10mM HEPES pH 7.6) and analyzed by live-cell imaging (described below) or fixed, permeabilized and mounted onto coverslips to analyze yeast MDC morphology in fixed cells. To this end, yeast cells with fixed for one hour at 30°C by addition of 4% (v/v) para-formaldehyde (Electron Microscopy Science) to the culture media. Fixed cells were isolated by centrifugation, washed with 100mM KPi (pH. 6.5) and incubated in DTT buffer (10mM 1,4-dithiothreitol (DTT), 100mM Tris, pH 9.4) for ten minutes. Subsequently, cell walls were digested for 30 minutes with 250 μg/ml zymolyase 100T (Amsbio; 120493-1) in zymolyase buffer (1.2M Sorbitol, 100mM KPi pH 7.4). Spheroplasts were re-isolated by centrifugation, washed with zymolyase buffer, loaded onto poly-L-Lysine coated multi-well slides (Thermo Fisher) and permeabilized with 100% methanol at −20°C for six minutes followed by a 30 second incubation in 100% acetone at −20°C. Slides were air-dried, mounted with ProLong Gold Antifade Mountant (Thermo Fisher) onto 1.5H coverslips according to manufacturer’s description and yeast cells were analyzed by microscopy.

### Mammalian Tissue Culture and Cell Lines

Immortalized *Miro1*^+/+^ and *Miro1*^-/-^ mouse embryonic fibroblasts (MEFs) were described previously (Schuler et al., 2017). *Miro2*^+/+^ and *Miro2*^-/-^ MEFs were isolated from *Miro2*^-/-^ and control littermates at embryonic gestation day 13 and immortalized by lentiviral transduction with simian virus 40 large T antigen as described previously (Nguyen et al., 2014; Zhu et al., 1991). *Miro2*^-/-^ mice were obtained from the mutant mouse resource and research Centers (MMRRC, UC Davis) and all experiments were approved by the University of Utah Institutional Animal Care and Use Committee. NIH 3T3 cells (CRL-1658), Cos-7 cells (CRL-1651) and HEK 293T cells (CRL-3216) were purchased from ATCC. The generation of MEFs stably expressing Tomm70A-EGFP and/or Timm50-mRFP is described below. All cell lines were maintained in high amino acid medium (DMEM high glucose, 4 mM L-Glutamine, no sodium pyruvate (Sigma Aldrich), 1x MEM non-essential amino acids (Sigma Aldrich), 10% (v/v) fetal bovine serum (FBS; Atlanta Biologicals)). Cell lines were routinely tested for mycoplasma contamination by PCR as previously described (Molla Kazemiha et al., 2009).

### Generation of Stable Cell Lines

MEFs stably expressing mTomm70A-EGFP and mTimm50-mRFP were generated by lentiviral transduction. To produce lentiviral particles, 293T cells were grown to 60% confluency and transfected with 3 μg packaging vector (pCMV-dR8.91), 0.3 μg envelope vector (pMD2.G) and 3 μg transfer vector (pLenti-mTomm70-EGFP or pLenti-mTimm50-mRFP). Medium was changed 18 hours after transfection and lentiviral particles were harvested 48 hours after transfection, sterile filtered and directly added for 24 hours to MEFs grown to 50% confluence. Five days post infection, stably transduced cells were with selected with 1mg/ml G418 and subsequently flow sorted for expression of Tomm70A-EGFP and/or Timm50-mRFP, respectively.

### Mammalian Transfections

To transiently express hSLC25A16-EGFP, TagBFP-C-KDEL or TagRFP-T-EEA1, MEFs were grown to 75% confluence and transfected with 500 ng plasmid per 100,000 cells using the Neon electroporation system (Invitrogen; 10 μl kit, one pulse, 1300 mV, 30 ms) according to manufacturer’s description. Transfected cells were directly plated onto glass coverslips or live-cell imaging dishes (Fisher Scientific) and grown for 24 hours in high amino acid medium prior to treatments.

### Mammalian MDC Assays

To visualize MDC formation in fixed mammalian cells, cells were plated at 50% confluence onto 1.5H glass coverslips (Fisher Scientific) in high amino acid medium and allowed to adhere for 24 hours. HEK 293T cells were plated onto L-polylysine (50 mg/ml for one hour in borate buffer) coated glass coverslips to promote cell adhesion. The next day, cells were treated with the indicated drug(s) in high amino acid medium, fixed and stained by indirect immunofluorescence for the indicated protein(s) and imaged as described below. Final drug concentrations used in mammalian MDC assays were 25 mg/ml cycloheximide (Sigma-Aldrich; C1988), 250 nM Torin1 (R&D Systems; 4247), 2 mM concanamycin A (Santa Cruz Biotechnology; sc-202111), 5 mg/ml tunicamycin (Sigma-Aldrich; T7765), 1 mM thapsigargin (Sigma-Aldrich; T9033) or 100 mM cobalt chloride (Sigma-Aldrich; C8661) for eight hours (unless otherwise indicated). Mitochondrial respiratory chain function was blocked with 10 mM rotenone (Sigma-Aldrich; R8875), 10 mM antimycin A (Sigma-Aldrich; A8674), 1 mM oligomycin (Sigma Aldrich; 75351) or 1 mM Carbonyl cyanide 4-(trifluoromethoxy)phenylhydrazone (FCCP) (Sigma-Aldrich; C2920). Glucose oxidase (Sigma-Aldrich; G2133) was used at 50 mU/ml to induce ROS formation. In amino acid starvation and amino acid re-addition experiments, cells were washed once with medium lacking all amino acids (EBSS, Invitrogen) and incubated in EBSS supplemented with the indicated amino acid (4 mM) (Fisher Scientific) for one hour prior to addition of 25 ug/ml cycloheximide to the medium.

### Indirect Immunofluorescence

To stain cells by indirect immunofluorescence, cells were grown on glass coverslips, treated as indicated, fixed with 4% paraformaldehyde (PFA; Electron Microscopy) in phosphate buffered saline (PBS) for 15 minutes at room temperature (RT) and permeabilized with 0.1% (v/v) Triton X-100 (BioRad) in PBS for 10 minutes at RT. Nonspecific antibody binding was blocked by incubating cells in 10% (v/v) normal goat serum (NGS; ThermoFisher) in PBS for one hour at RT. The indicated primary antibody (Tomm70A (Sigma HPA048020, 1:400 and Santa Cruz sc-390545, 1:200), Tomm20 (Millipore MABT116, 1:400), Cytochrome c (BD Bioscience 556432, 1:200), AtpB (ABCAM ab14730, 1:400), PDH E2/3 (ABCAM ab110333, 1:200), Prx3 (ABCAM ab73349, 1:200), Pex14 (Millipore ABC142, 1:200) was applied in PBS + 10% NGS for 1 hour at RT. Primary antibodies were detected by incubating cells for 30 minutes with AlexaFluor conjugated secondary antibodies made in goat (ThermoFisher, 1:200 in PBS + 10% NGS). Coverslips were mounted onto microscopy slides (Fisher Scientific) using Prolong Diamond Antifade (ThermoFisher) and were cured for 24 hours at RT prior to imaging.

### Mammalian Live-Cell Imaging

To visualize MDCs in live cells, MEFs stably expressing either mTomm70A-EGFP alone or mTomm70A-EGFP and mTimm50-mRFP were plated onto live-cell imaging dishes (CellVis) and grown for 24 hours in live-cell imaging medium (Fluorobrite (Invitrogen), 4 mM L-Glutamine, (Sigma Aldrich), 1x MEM non-essential amino acids (Sigma Aldrich), 10% (v/v) FBS). The next day, cells were treated as indicated in live-cell imaging medium and 185 nm optical Z-sections were acquired with an Airyscan LSM880 as described below. Time-lapse images and movies were acquired at 15 second intervals with an Airyscan LSM880 in Airyscan Fast mode over 60 - 120 minutes and show maximum intensity projections generated in Fiji (Schindelin et al., 2012).

The mitochondrial inner membrane potential was visualized by staining cells with 50 nM tetramethylrhodamine methyl ester (TMRM) for 10 minutes in live-cell imaging medium and lysosomes were visualized by staining cells with 50nM Lysotracker Red DND-99 (Thermo Fisher) for 30 minutes in live-cell imaging medium according to manufacturer’s description. After staining, media were replaced with dye-free live-cell imaging media. To visualize the ER and endosomes, cells were transfected with pTagBFP-C-KDEL or TagRFP-T-EEA1 prior to plating onto live-cell imaging dishes.

### Microscopy and Image analysis

200 nm optical Z-sections of live or fixed yeast and mammalian cells were acquired with an AxioImager M2 (Carl Zeiss) equipped with an Axiocam 506 monochromatic camera (Carl Zeiss) 63× and 100× oil-immersion objective (Carl Zeiss, Plan Apochromat, NA 1.4), or, for super-resolution images, with an Airyscan LSM800 or LSM880 (Carl Zeiss) equipped with an Airyscan detector (Carl Zeiss) and 63× oil-immersion objective (Carl Zeiss, Plan Apochromat, NA 1.4). All images were acquired with ZEN (Carl Zeiss) and super-resolution images were further processed using the automated Airyscan processing algorithm in ZEN (Carl Zeiss). Individual channels of all images were minimally adjusted in Fiji to match the fluorescence intensities between channels for better visualization. All images were cropped in Fiji or Photoshop CC (Adobe) and annotated in Illustrator (Adobe). A single focal plane is displayed for yeast images in Figure 1A and all images of live-mammalian cells. All images of fixed (and stained) yeast and mammalian cells show maximum intensity projections generated in Fiji and cropped in Adobe Photoshop CC.

Line-scan analysis and measurements of mammalian MDC size were performed on non-adjusted, single Z-sections from super resolution images. To quantify the average MDC size, the diameter of spherical MDCs that had a visible lumen was measured using the line tool in Fiji. Tubular MDCs or smaller Tomm70A-enriched structures lacking Timm50-mRFP were excluded from the analysis since they likely represent growing MDCs. The number of MDCs per cell and the average frequency of cells with at least one MDC were quantified throughout the entire Z-series of the acquired data. For quantification of MDC formation in HEK 293T cells, the raw data was processed by iterative deconvolution in ZEN (Carl Zeiss) to better resolve the dense mitochondrial network in these cells. Mammalian MDCs were identified as large, round, Tomm70A-containing structures that were brighter than the average mitochondrial tubule, had at least the diameter of a mitochondrion and were not stained with the mitochondrial matrix marker AtpB (for fixed cells) or the inner membrane marker Timm50-mRFP (for live cells). Smaller, less bright or irregularly shaped Tomm70A puncta were excluded from the quantification.

### Protein Preparation and Immunoblotting

For western blot analysis, cells were grown to 80% confluency on 6-well plates and treated as indicated. To prepare whole cell lysates, cells were washed with tris buffered saline (TBS) and lysed in 250 μl radioimmunoprecipitation assay (RIPA) buffer (25 Tris pH 8, 150 mM NaCl, 1 mM EDTA, 0.1% [w/v] sodium dodecyl sulfate (SDS), 0.1% [w/v] sodium deoxycholate, 1% [v/v] Triton X-100) containing 1× Protease/Phosphatase inhibitor (Pierce) for 15’ on ice. Cell lysates were cleared by centrifugation, protein concentrations were determined by a bicinchoninic assay (G Biosciences) according to manufacturer’s instructions and equal amounts of protein were denatured for 5 minutes at 95°C with Laemmli buffer (63 mM Tris pH 6.8, 2% (w/v) SDS, 10% (v/v) glycerol, 1 mg/ml bromophenol blue, 1% (v/v) β-mercaptoethanol). For analysis of the Prx2/3 monomer-to-dimer ratio, 50 mM N-ethylmaleimide (Sigma Aldrich) was added to the RIPA buffer and no β-mercaptoethanol was added during the denaturation step. Samples were subjected to SDS polyacrylamide gel electrophoresis and transferred to PVDF membrane (Millipore) by wet transfer. Unspecific antibody binding was blocked by incubation with TBS containing 5% (w/v) bovine serum albumin (Sigma Aldrich) for one hour at RT. After primary antibody (S6-Kinase phospho Thr 389 (CST 9234, 1:1000), S6-Kinase (CST 2708, 1:1000), 4EBP (CST 9644, 1:1000), GAPDH (Thermo Fisher PA1-987, WB: 1:1000), Prx2 (ABCAM ab109367, 1:1000), Prx3 (ABCAM ab73349, 1:1000)) incubation overnight at 4°C, membranes were washed five times with TBS and incubated with secondary antibody (goat-anti-rabbit HRP-conjugated,1:2000 in TBS + 5% dry milk, Sigma Aldrich) for one hour at RT. Membranes were washed five times with TBS, enhanced chemiluminescence solution (Thermo Fisher) was applied and the antibody signal was detected with a BioRad Chemidoc MP system. All blots were exported as TIFFs and cropped in Adobe Photoshop CC. Western blots show one representative blot from *n* = 3 replicates performed in parallel with the associated MDC assay.

### Whole-Cell Metabolite Analysis

For analysis of whole cell metabolite levels in mammalian cells, MEFs were grown to 80% confluency on 150 mm tissue culture dishes and treated with 25 μg/ml cycloheximide or DMSO for four hours. Cells were washed once, scraped into ice cold PBS, centrifuged at 100 ×g for three minutes in microcentrifuge tubes, and cell pellets were shock frozen in liquid nitrogen. 90% cold methanol solution containing the internal standard succinic-*d*4 acid was added to each cell pellet and samples were incubated for one hour at −20°C. Subsequently, cell debris were removed by centrifugation and the supernatant was dried *en vacuo*. Amino acid composition analysis was performed on an Agilent 7200 GC-QToF mass spectrometer fitted with an Agilent 7890 gas chromatograph and an Agilent 7693A automatic liquid sampler. Dried samples were suspended in 40 μl of 40 mg/ml O-methoxylamine hydrochloride (MP Biomedicals) in dry pyridine and incubated for one hour at 30°C. 25 μl of this solution was added to auto sampler vials. 60 μl of N-methyl-N-trimethylsilyltrifluoracetamide (Pierce) was added automatically via the auto sampler and incubated for 30 minutes at 37°C with shaking. After incubation, 1 μl of the prepared sample was injected into the gas chromatograph inlet in the split mode with the inlet temperature held at 250°C. A 25:1 split ratio was used for analysis. The gas chromatograph had an initial temperature of 60°C for one minute followed by a 10°C/min ramp to 325°C and a hold time of 2 minutes. A 30-meter Agilent Zorbax DB-5MS with 10m Duraguard capillary column was employed for chromatographic separation. Helium was used as the carrier gas at a rate of 1 ml/min. Data was collected using MassHunter software (Agilent). Metabolites were identified and their peak area was recorded using MassHunter Quant. Metabolite identity was established using a combination of an in-house metabolite library developed using pure purchased standards, the NIST library and the Fiehn Library. Values for each metabolite were normalized to the total ion count and the internal standard in each sample and are displayed as fold change to DMSO. Error bars show the mean ± standard error from three biological replicates analyzed in the same GC-MS run.

### Statistics

Experiments were repeated at least three times. All attempts at replication were successful. Sample sizes were as large as possible to be representative, but of a manageable size to quantify MDCs. Unless indicated, quantifications show the mean ± standard error from three biological replicates with *n* = 50 cells per experiment. No data were excluded from the analyses. No randomization or blinding was used as all experiments were performed with defined laboratory reagents, yeast strains of known genotypes, and mammalian cell lines purchased from ATCC or generated and characterized in house.

### Data Availability

The data that support the findings of this study are available from the corresponding author upon request.

---

**Figure 1 Supplement 1.**
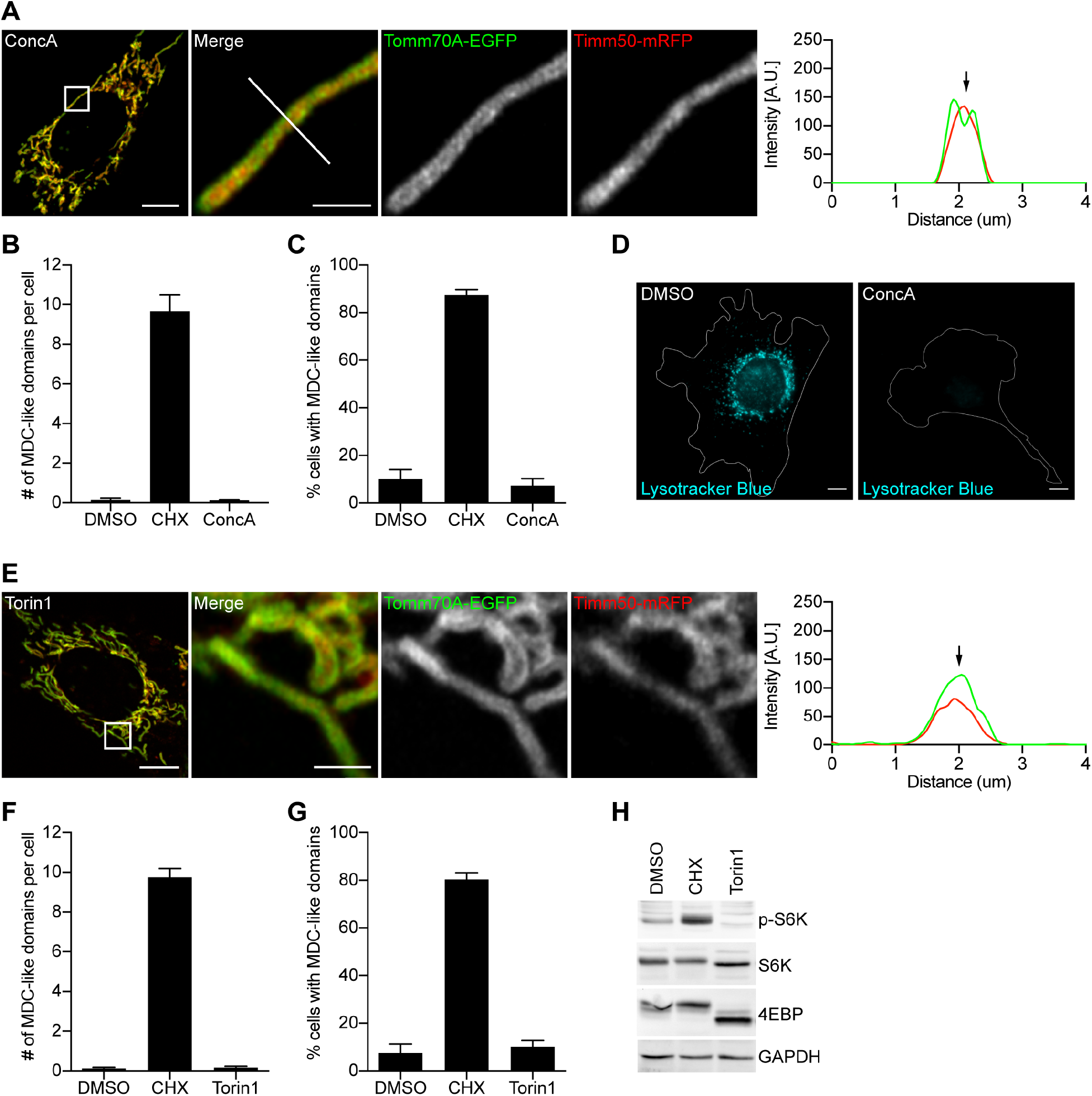
V-ATPase and mTOR inhibition do not activate formation of MDC-like domains in mammals. (A) Images of concanamycin A (ConcA) treated MEFs expressing Tomm70A-EGFP and Timm50-mRFP. White line marks fluorescence intensity profile position. Black arrow marks mitochondrial tubule on line-scan analysis plot. Scale bars, 10 μm and 2 μm. (B, C) Quantification of MDC-like domain formation in MEFs treated with DMSO, cycloheximide (CHX) or ConcA. Error bars show mean ± SE from three replicates with *n* = 50 cells per replicate. (D) Images of DMSO or ConcA treated MEFs stained with pH-sensitive Lysotracker Blue. Scale bar, 10 μm. (E) Images of Torin1 treated MEFs expressing Tomm70A-EGFP and Timm50-mRFP. White line marks fluorescence intensity profile position. Black arrow marks mitochondrial tubule on line-scan analysis plot. Scale bars, 10 μm and 2 μm. (F, G) Quantification of MDC-like domain formation in MEFs treated with DMSO, CHX or Torin1. Error bars show mean ± SE from three replicates with *n* = 50 cells per replicate. (H) Western Blot showing phosphorylation of mTOR substrates in MEFs. GAPDH, loading control. (B, F) Number of MDC-like domains per cell. (C, G) Percentage of cells with at least one MDC-like domain.

**Figure 1 Supplement 2.**
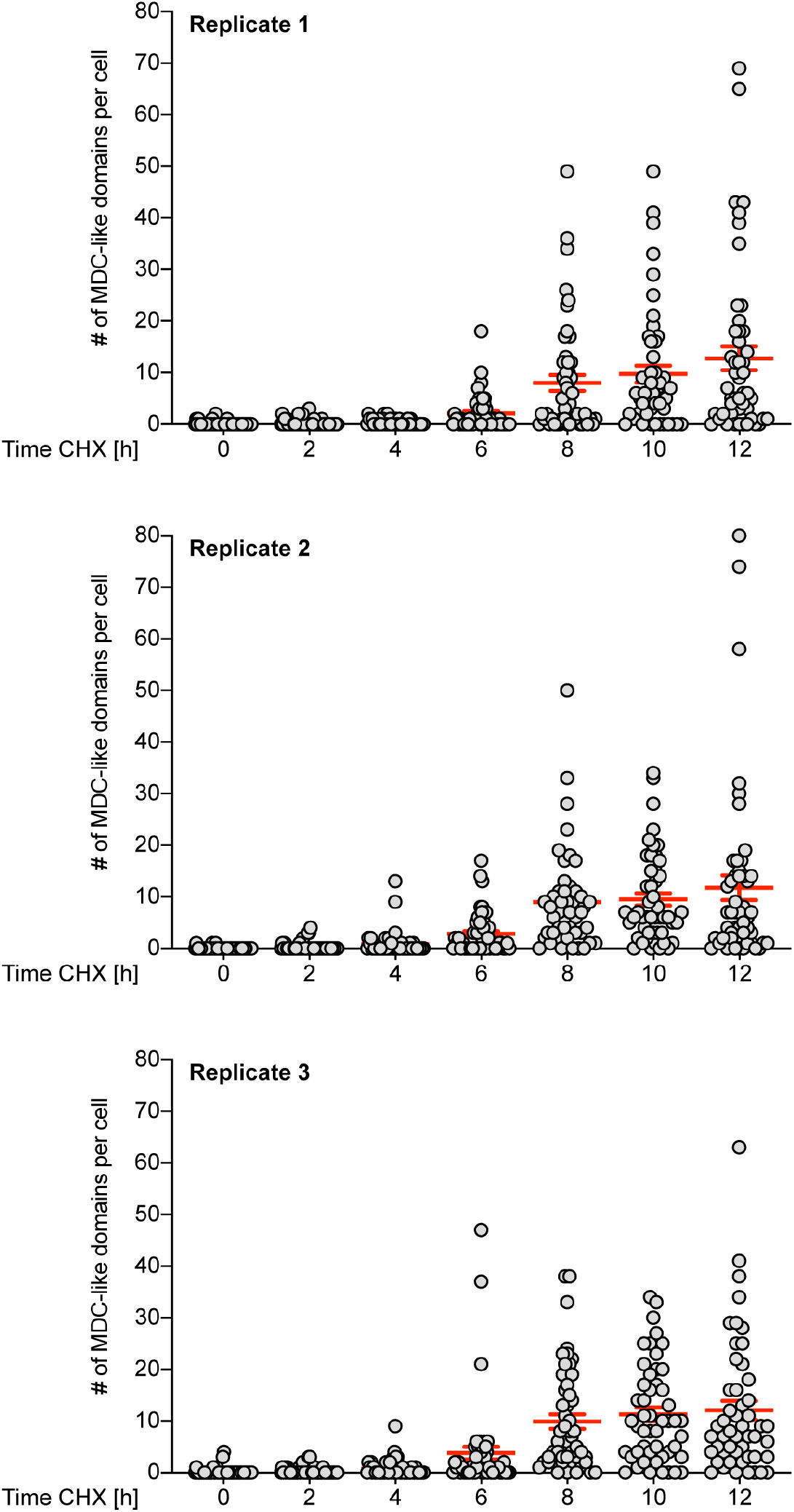
Mammalian cells form multiple MDC-like domains. Scatter plots showing the number of cycloheximide (CHX)-induced MDC-like domains per cell over time in *n* = three independent replicates. Error bars show mean ± SD of *n* = 50 cells per replicate and time point.

**Figure 2 Supplement 1.**
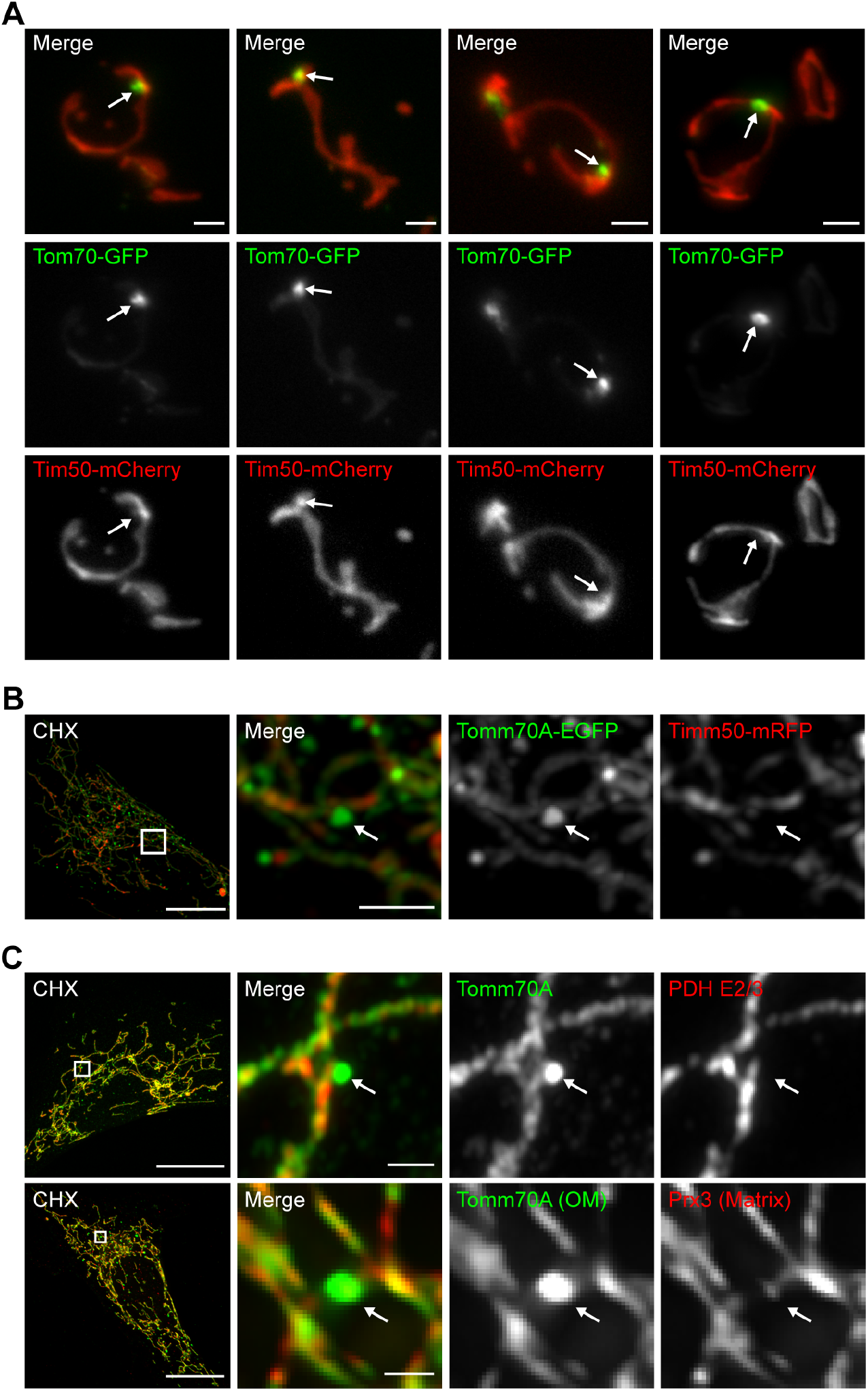
Yeast MDCs and mammalian MDC-like domains are sensitive to fixation. (A) Images of cycloheximide (CHX)-induced MDCs in Tom70-GFP and Tim50-mCherry expressing yeast cells after fixation and permeabilization. White arrow marks MDC. Scale bar, 2 μm. (B) Images of cycloheximide (CHX)-induced MDC-like domains in MEFs expressing Tomm70A-EGFP and Timm50-mRFP after fixation and permeabilization. (C) Images of CHX treated MEFs, fixed and stained for endogenous Tomm70A and the indicated mitochondrial matrix protein. (B, C) White arrow marks MDC-like domain. Scale bars, 10 μm and 2 μm.

**Figure 2 Supplement 2.**
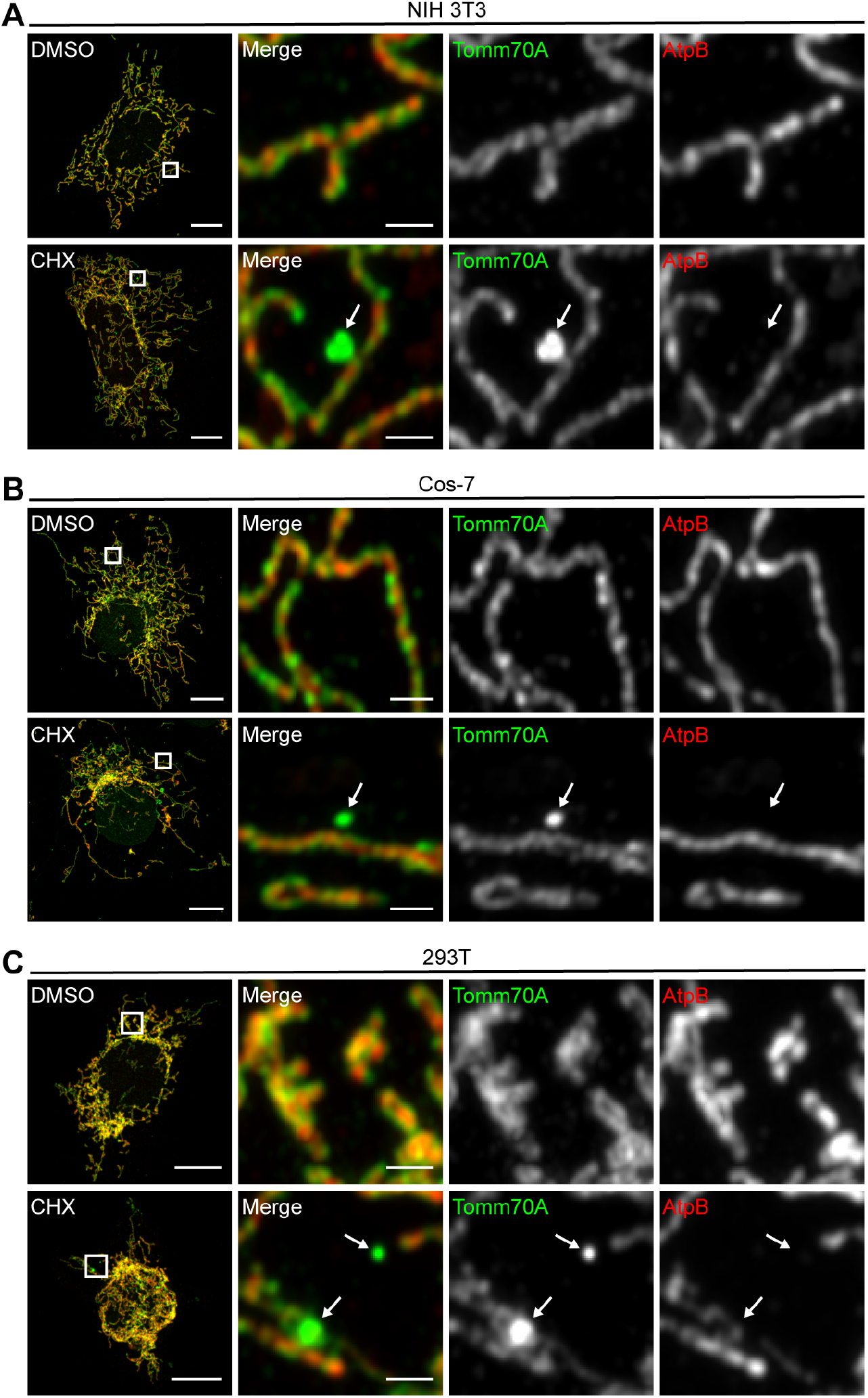
Translation inhibition activates formation of MDC-like domains in numerous cell lines. (A – C) Images of DMSO or CHX treated (A) NIH 3T3 cells, (B) Cos-7 cells and (C) 293T cells, fixed and stained for endogenous Tomm70A and the mitochondrial matrix protein AtpB. White arrow marks MDC-like domain. Scale bars, 10 μm and 2 μm.

**Figure 4 Supplement 1.**
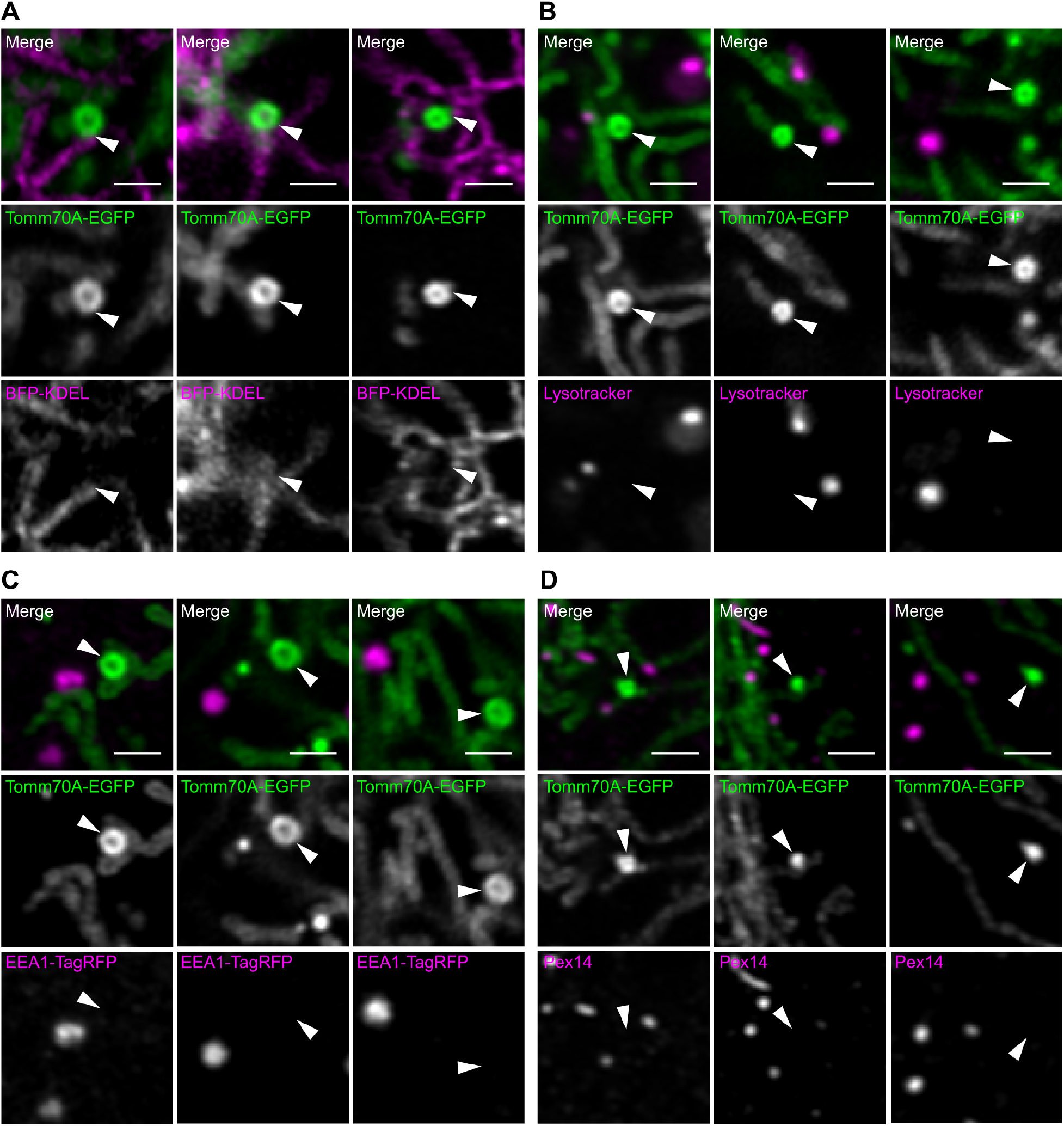
Association of mammalian MDC-like domains with the ER, lysosomes, endosomes and peroxisomes. (A) Images of cycloheximide (CHX)-induced MDC-like domains in MEFs expressing Tomm70-A-EGFP and the ER marker BFP-KDEL. (B) Images of CHX-induced MDC-like domains in MEFs expressing Tomm70-A-EGFP and stained with the pH-sensitive dye Lysotracker Red. (C) Images of CHX-induced MDC-like domains in MEFs expressing Tomm70-A-EGFP and the endosomal protein EEA1 fused to TagRFP. (D) Images of CHX-induced MDC-like domains in MEFs expressing Tomm70-A-EGFP, fixed and stained for peroxisomal protein Pex14. (A – D) Arrowhead marks MDC-like domain. Scale bar, 2 μm.

**Figure 6 Supplement 1.**
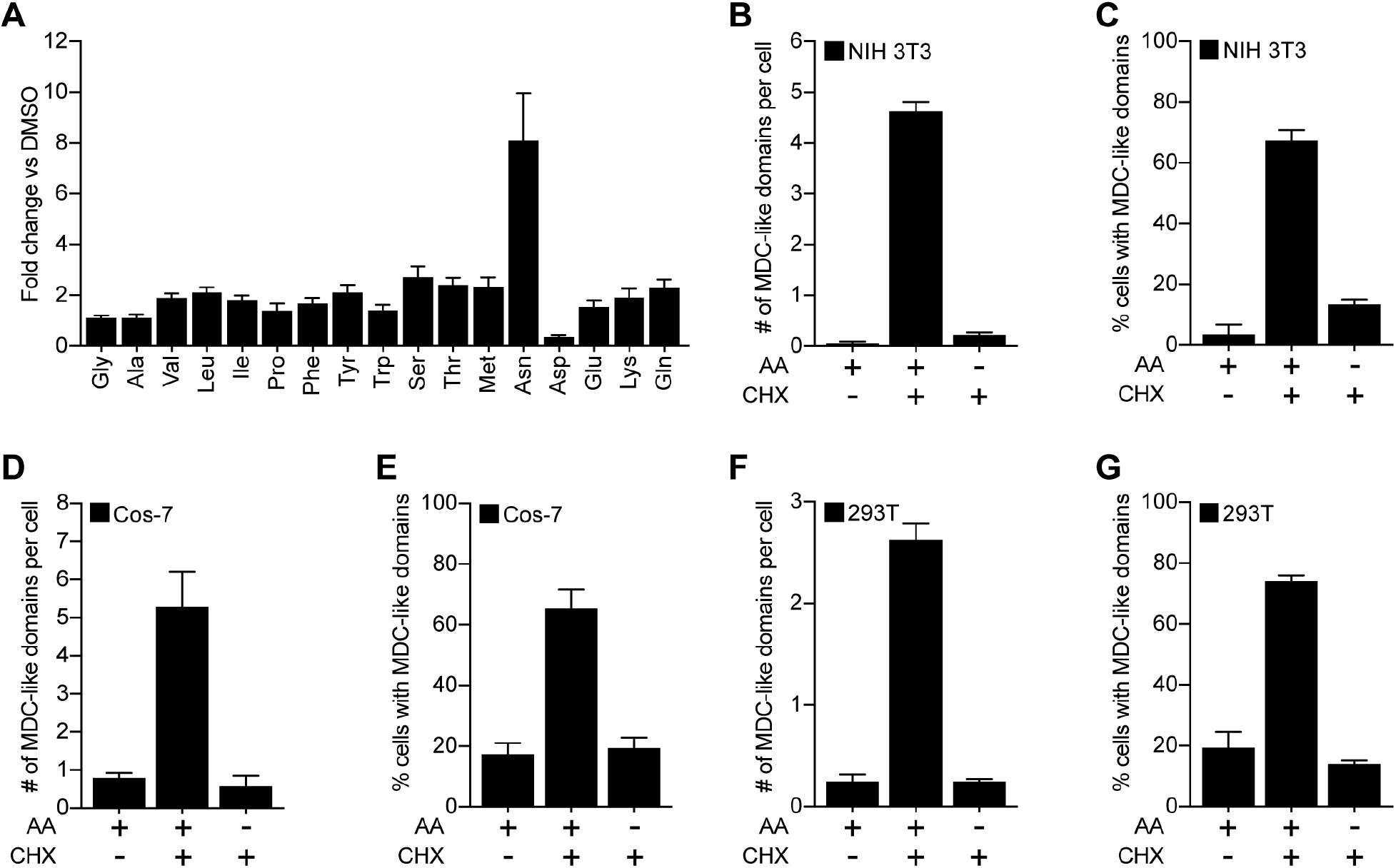
Regulation of MDC-like domain formation by amino acids is conserved across cell lines. (A) Whole-cell amino acid levels in cells treated with cycloheximide (CHX) compared to DMSO treated controls. *N* = 5 biological replicates. (B – G) Quantification of CHX-induced MDC-like domain formation in (B, C) NIH 3T3 cells, (D, E) Cos-7 cells and (F, G) 293T cells in presence or absence of external amino acids. Error bars show mean ± SE from three replicates with *n* = 50 cells per replicate. (B, D, F) Number of MDC-like domains per cell. (C, E, G) Percentage of cells with at least one MDC-like domain.

**Figure 6 Supplement 2.**
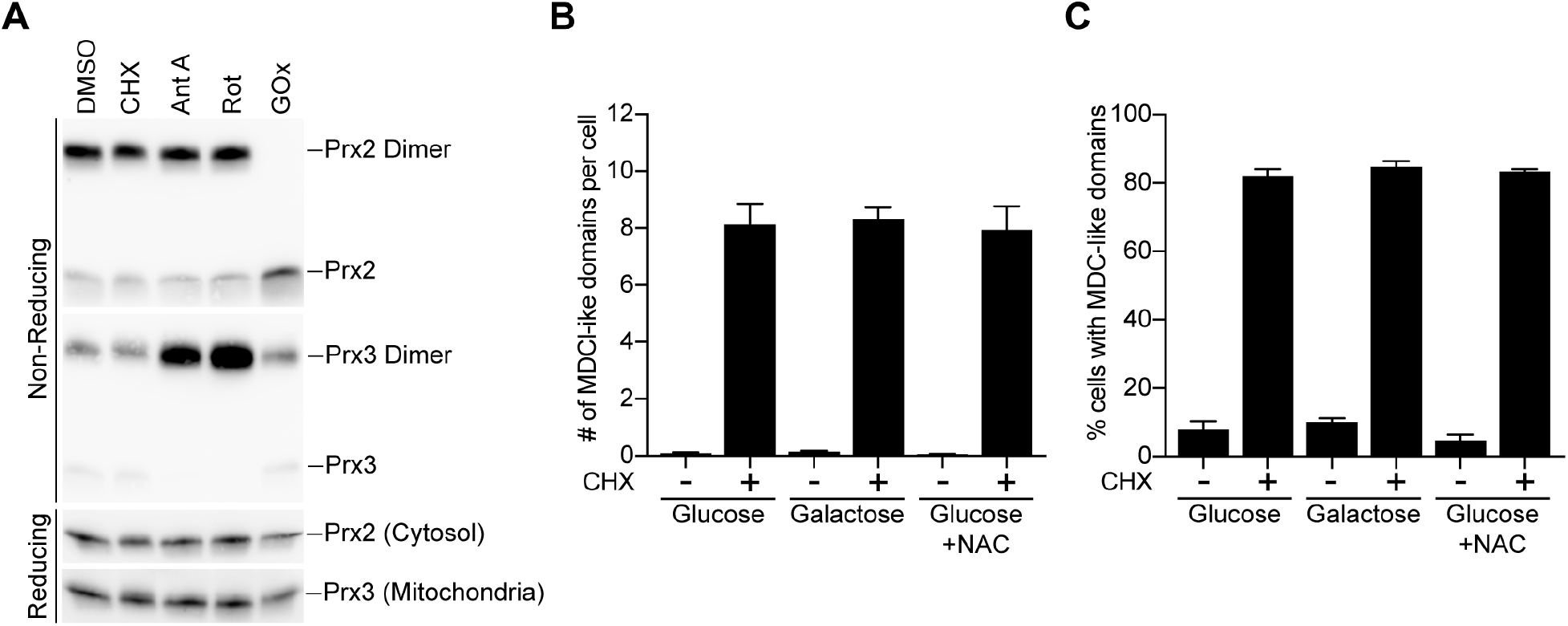
ROS stress does not activate formation of MDC-like domains in mammalian cells. (A) Reducing and non-reducing SDS-PAGE and Western blot analysis of monomer to dimer ratio of cytosolic (Prx2) and mitochondrial (Prx3) peroxiredoxin in MEFs treated with DMSO, cycloheximide (CHX), antimycin A (AntA), rotenone (Rot) or glucose oxidase (GOx). A shift in the monomer to dimer ratio indicates a change in the subcellular redox status. (B, C) Quantification of CHX-induced MDC-like domain formation in the indicated media. N-acetylcysteine, NAC. Error bars show mean ± SE from three replicates with *n* = 50 cells per replicate. (B) Number of MDC-like domains per cell. (C) Percentage of cells with at least one MDC-like domain.

**Movie 1. Related to Figure 3A.** Movie showing cycloheximide-induced formation of MDC-like domains in mammalian cells. MEFs expressing Tomm70A-EGFP were pre-treated with cycloheximide for six hours and imaged every 15 seconds for two hours. Formation of MDC-like domains begins with widening of the mitochondrial network followed by concentration of Tomm70A-GFP into the growing MDC and recovery of mitochondrial network morphology. Upon formation, the MDC-like domain remains associated with the mitochondrial network.

**Movie 2. Related to Figure 3B.** Movie showing cycloheximide-induced formation of MDC-like domains in mammalian cells. MEFs expressing Tomm70A-EGFP were pre-treated with cycloheximide for six hours and imaged every 15 seconds for two hours. Formation of MDC-like domains is preceded by widening of the mitochondrial network followed by concentration of Tomm70A-GFP into the growing MDC and recovery of mitochondrial network morphology. Note that the newly forming and preformed MDC-like domains stably associate with the mitochondrial network.

**Movie 3. Related to Figure 3C.** Movie showing dynamics of MDC-like domains in mammalian cells. MEFs expressing Tomm70A-EGFP were treated with cycloheximide for seven hours and imaged every 15 seconds for one hour. MDC-like domains dynamically interact with the local mitochondrial network. These events include release from and docking to mitochondria, as well as movement along the mitochondrial tubule.

